# Crystal structures reveal that Lewis-x and fucose bind to secondary cholera toxin binding site – in contrast to fucosyl-GM1

**DOI:** 10.1101/431130

**Authors:** Joel B. Heim, Vesna Hodnik, Julie E. Heggelund, Gregor Anderluh, Ute Krengel

## Abstract

Cholera is a life-threatening diarrhoeal disease caused by the human pathogen *Vibrio cholerae*. Infection occurs after ingestion of the bacteria, which colonize the human small intestine and secrete their major virulence factor - the cholera toxin (CT). Recent studies suggest that the GM1 receptor may not be the only target of the CT, and that fucosylated receptors such as Lewis^x^ (Le^x^) and histo-blood group antigens may also be important for cellular uptake and toxicity. However, where and how Le^x^ binds to the CT remains unclear. Here we report the high-resolution crystal structure (1.5 Å) of the receptor-binding B-subunit of the CT bound to the Le^x^ trisaccharide, and present matching SPR data for CT holotoxins. Le^x^, and also L-fucose alone (at 500-fold molar excess), bind to the secondary binding site of the toxin, distinct from the GM1 binding site. In contrast, fucosyl-GM1 mainly binds to the primary binding site due to high-affinity interactions of its GM1 core. The two binding sites are likely connected by allosteric cross-talk, potentially affecting toxin uptake. We also discuss why secretors are protected from severe cholera.

**Author summary:** Cholera is a severe diarrhoeal disease that is still a major killer in many parts of the world, especially in regions struck by natural disasters and wars. However, some individuals experience milder cholera symptoms. These so-called ‘secretors’, who have blood group antigens also in their bodily fluids like their saliva and the slimy mucus layer covering their stomach and intestines, appear to be somewhat protected. Here we present detailed atomic structures of cholera toxin and quantitative binding data that give clues of the protective mechanisms. Interactions of the protein toxin with sugar molecules are of crucial importance both for toxicity and protection. In addition, we identify a new tool for biochemical studies, and lay the groundwork for the design of cholera drugs and vaccines that may save countless human lives.

## Introduction

Cholera is an acute and severe diarrhoeal disease that if left untreated can cause serious dehydration and death within hours^1^. The disease is a major threat, especially in countries lacking proper sanitation, in war zones and after natural disasters. The current outbreak in Yemen (2016-2018) is the worst in recent history with more than one million reported cases and more than 2,200 deaths^2,3^.

Cholera is easily transmitted between humans through the faecal-oral route, *e.g*. by consuming food or water contaminated with the bacterium *Vibrio cholerae*. The bacteria colonize the small intestine and secrete the cholera toxin (CT)^4^. CT belongs to the protein family of AB_5_ toxins that consist of one catalytically active A-subunit and a homopentamer of non-toxic B-subunits^4,5^. The CT B-pentamer (CTB) facilitates binding to epithelial cells in the small intestine and subsequent cellular uptake of the holotoxin^4, 6^. In the cytosol, the A-subunit hijacks the cells’ own signalling pathways, causing the secretion of chloride ions into the intestinal lumen and, due to the osmotic gradient, ultimately the severe watery diarrhoea typical of cholera^6–8^.

Key aspects of cholera intoxication are still poorly understood, starting with the interplay between the toxins and their cellular receptors, and the cellular uptake mechanism. For decades, the GM1 ganglioside has been considered to be the only cellular receptor for the CT^9–11^. The binding of CT to the branched GM1 pentasaccharide is one of the strongest protein-carbohydrate interactions known (*K*_d_=43 nM)^12^ and is mainly due to the two terminal sugar residues galactose and sialic acid, which bind the toxin with a “two-fingered grip”^13^. A large body of research points towards this high affinity interaction being the main entry pathway for the CT^10,14–18^. For example, the incorporation of exogenously added GM1 in human intestinal mucosal cells or rabbit small bowel segments lead to a concentration-dependent increase in the number of toxin receptors and in the sensitivity to the diarrheogenic action of CT^10, 14^. CT binds GM1 at the bottom surface of the B-pentamer, which serves as a landing platform on the cell membrane^13^. More recently, CT was found to bind histo-blood group antigens (BGAs) at a secondary binding site on the lateral side of the B-pentamer^19^, which had originally been discovered for chimeras of CTB and the homologous *Escherichia coli* heat-labile enterotoxin (LTB)^20^. Cholera toxin naturally occurs in two variants - classical CT (cCT) and El Tor CT (ET CT), which are produced by the corresponding *V. cholerae* biotypes^4^. However, since 2001 a new vibrio strain has been observed with characteristics of the El Tor biotype that produces CT with the classical amino acid sequence^21^. The two variants only differ in residues 18 and 47 in the secondary CT binding site (cCT: H18, T47, ET CT: Y18, I47)^4, 22^, which is why they have been in the focus of cholera blood group binding studies^19,23-25^. Moreover, there is increasing evidence that GM1 is not the only CT receptor and may not even be its main receptor^11,20,26–33^.

Current research shows that fucosylated structures have an important role in cholera intoxication^30–33^. CT binds fucosylated human BGAs merely with millimolar affinity^19,23–25^, but nevertheless causes blood-group-dependent cellular intoxication^31^. Patients with blood group O are also more likely to get severe cholera symptoms^34–37^. BGAs are present in the intestine in high concentrations, especially compared to GM1^38^. Most recently, Le^x^, a BGA precursor expressed on human granulocytes and intestinal cells, has been shown to specifically bind to CTB and was proposed to function as a cellular receptor^32^. Cholera intoxication in mice can be independent of GM1^32^, and fucose-based inhibitors, especially a fucose-containing polymer, are potent inhibitors of CT binding to intestinal cells^33^. It is, however, under debate if and how Le^x^ and related structures bind to the GM1 binding site, the BGA binding site or both^32^. Likewise, it is unknown where and how L-fucose binds to the CT.

Here we investigated the binding of CTB to the Le^x^ trisaccharide, L-fucose and the fucosyl-GMl oligosaccharide (os) (Figure 1) by X-ray crystallography and surface plasmon resonance (SPR) spectroscopy. We hypothesized that Le^x^ and related sugars interact solely with the secondary binding site of CTB based on their similarity to known toxin-bound BGAs and human milk oligosaccharides (HMOs)^19^. To test this hypothesis, we crystallized CTB with the Le^x^ trisaccharide or L-fucose. Indeed we found both sugars in the secondary, but not in the primary binding sites. In contrast, fucosyl-GM1os occupied the primary GM1 receptor site (and some of the secondary binding sites, *via* its fucose), as expected due to the similarity of the two compounds. It binds the CT with the same strong affinity as GM1 and with a two-fingered grip, but the fucose residue contributes little to their interaction, serving more as decoration.

**Figure 1.**
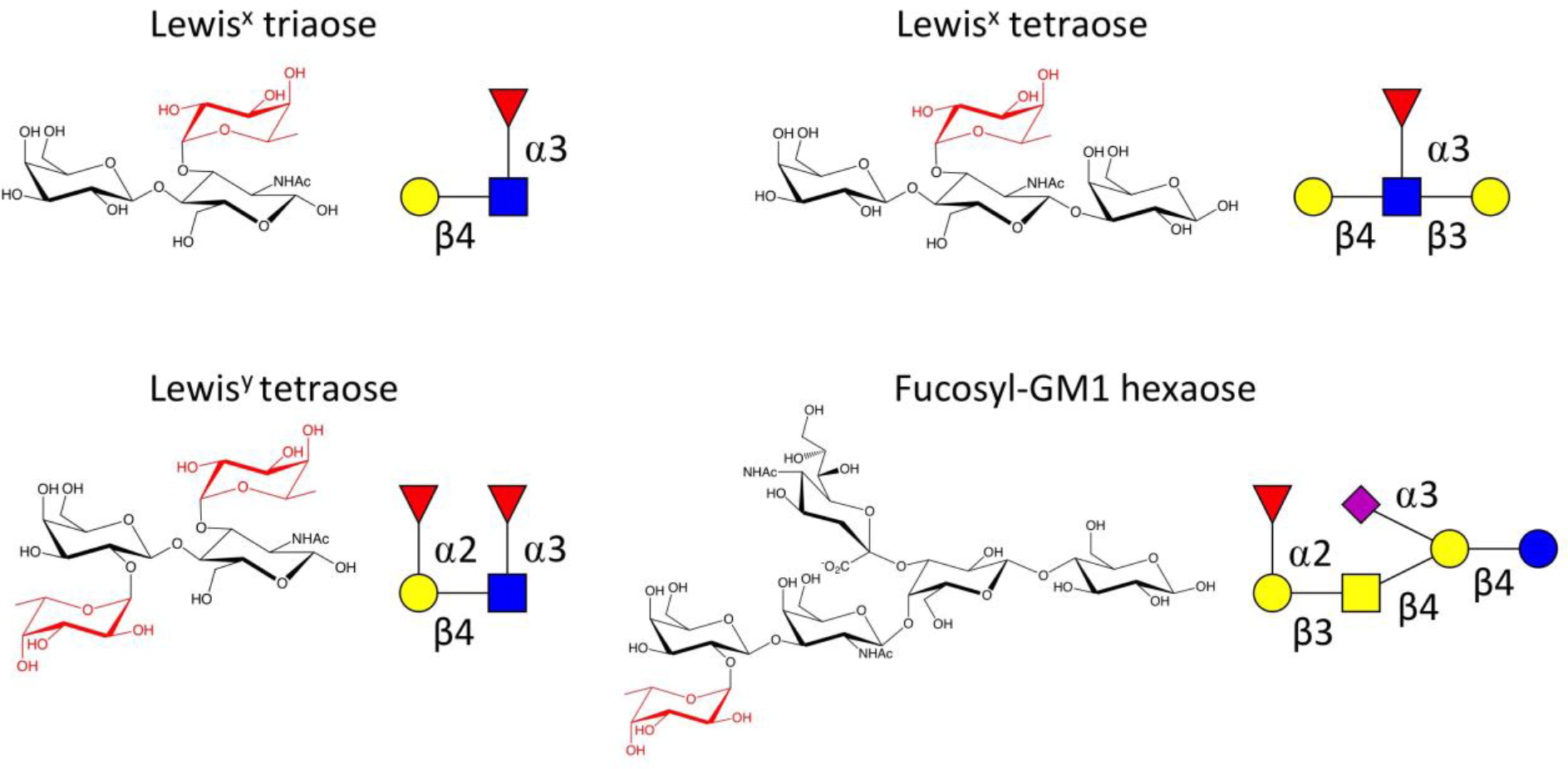
Structures of the oligosaccharides Lewis^x^ triaose, Lewis^x^ tetraose, Lewis^y^ and fucosyl-GM1 hexaose. Fucose residues are highlighted in red. Carbohydrate symbols follow the nomenclature of the Consortium for Functional Glycomics (Nomenclature Committee, Consortium for Functional Glycomics; D-galactose (Gal)-yellow circle, *N*-acetylgalactosamine (GalNAc)-yellow square, D-glucose (Glc)-blue circle, *N*-acetylglucosamine (GlcNAc)-blue square, L-fucose (Fuc)-red triangle.

## Results

### Crystal structures

#### CTB in complex with Le^X^

To study the interaction of CT with Le^x^, we crystallized purified classical CTB (cCTB) in complex with Le^x^ triaose. Crystals were obtained in two different crystallization conditions. They belonged to space group *P*2_1_2_1_2_1_ and contained two CTB pentamers in the asymmetric unit. Therefore, we can observe 20 crystallographically distinct primary binding and secondary binding sites. The protein structures show the typical “doughnut-shaped” CTB structure of five symmetrically arranged B subunits, each consisting of two three-stranded antiparallel β-beta-sheets with a-helices on both sides^13^. The structures were determined to 2.0 Å and 1.5 Å resolution, respectively. In both crystal forms, Le^x^ is only observed in the secondary binding sites of the CT. The binding mode and interactions of Le^x^ described here are based on the 1.5 Å resolution structure, which was refined to a high quality model (Figure 2 a, Table 1; *R*_free_ = 22.4%). Generally, the structure is well defined except for the flexible loop comprising residues 50-61 and the C-terminal asparagine residue, which exhibits some disorder. Inspection of the electron density maps revealed the presence of Le^x^ triaose in eight of ten secondary binding (Figure 2 b). One additional binding site contained electron density in sufficient quality to place the terminal L-fucose, and the last secondary binding site was blocked by crystal contacts. No ligand density was observed in any of the primary binding sites, even at low sigma cut-offs.

**Figure 2.**
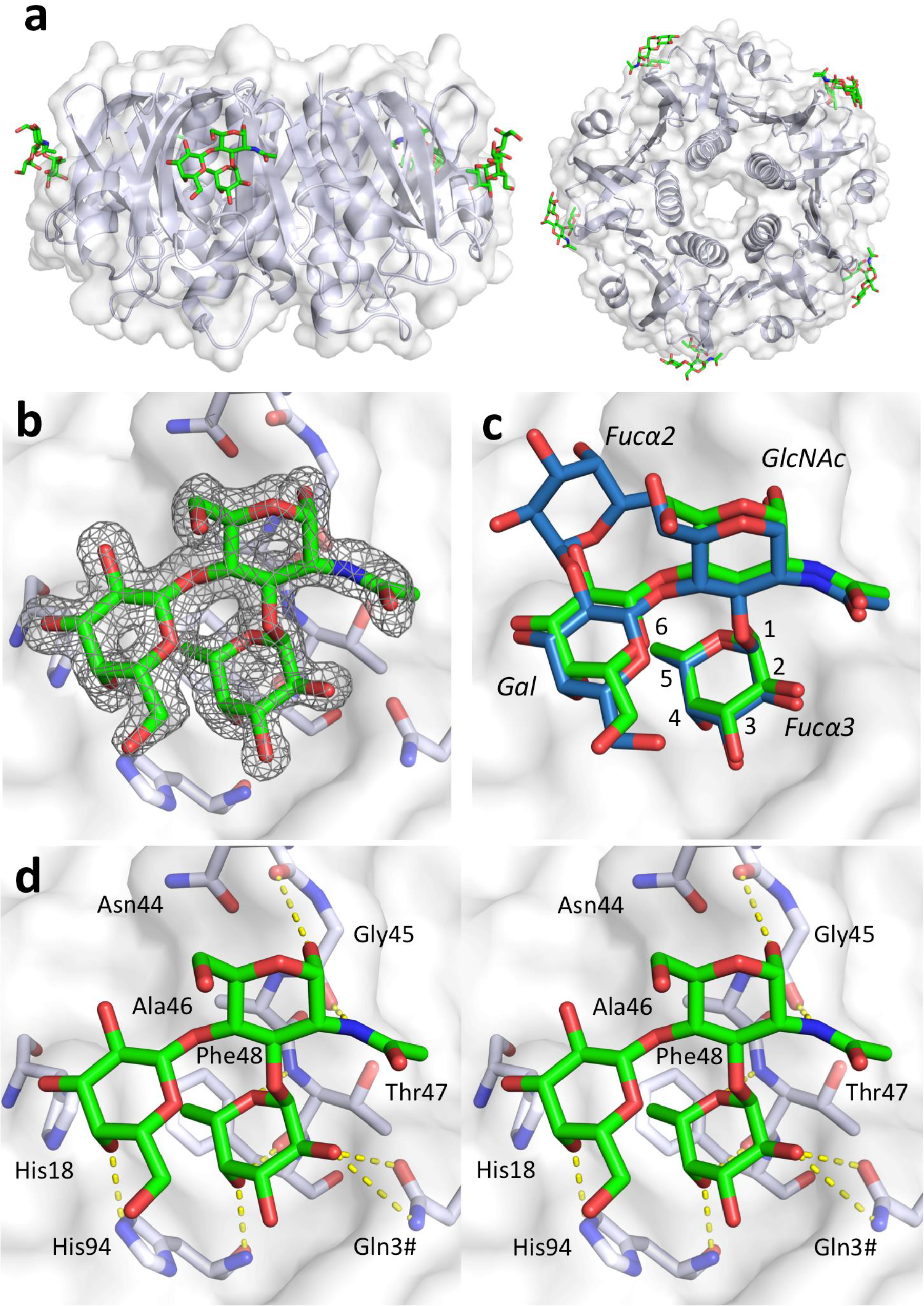
Lewis^x^ binds to the secondary binding site of the cholera toxin. (a) X-ray structure of cCTB in complex with Le^x^ (PDB ID: 6HJD, this work); side and top views (rotated by 90°). The toxin B-pentamer is shown in cartoon (light blue) and transparent surface representation (white), and the ligands depicted in stick representation (green). Le^x^ only binds to secondary binding sites on the lateral side of the toxin. (b) Close-up view of the secondary binding site (chain B), with σA-weighted F_o_-F_c_ electron density map for Le^x^ (grey mesh, contoured at 3.0 σ, generated before placing the ligand) and selected residues depicted in stick representation. (c) Superimposition of Le^y^ tetraose (blue sticks; PDB ID: 5ELB^19^) on cCTB complex with Le^x^ triaose (green sticks; PDB ID: 6HJD, this work). Carbohydrate residues are labelled in italics, and fucose carbons are numbered. (d) Stereo image of the carbohydrate-toxin interactions. Hydrogen bonds are shown as yellow dashed lines, amino acid residues are labelled with 3-letter code. The figure was prepared with MacPyMol (Schrödinger LLC (https://www.schrodinger.com/pymol); version 1.8.0.3).

Le^x^ triaose superimposes well with the trisaccharide core of the previously reported Lewis antigens^19^ (Figure 2 c), and engages in similar H-bonding interactions (Figure 2 d, Supplementary Table S1)^19^. Its reducing end (*N*-acetylglucosamine; GlcNAc) points towards residue 47. The blood-group antigen binding site of CT targets Lewis antigens mainly through the α1,3 fucose (although Le^y^ itself can bind the CT in two orientations, where either Fucα2 or Fucα3 can serve as main interaction points). As observed for cCTB bound to Le^y^ and A-Le^y^ (5ELB, 5ELD^19^), the fucose forms hydrogen bonds to Gln3# from the adjacent B-subunit and to the backbones of residues 47 and 94 (Figure 2 d, Supplementary Table S1). These data confirm our hypothesis that Le^x^ binds to the secondary binding site, similarly to BGAs.

**Table 1.**
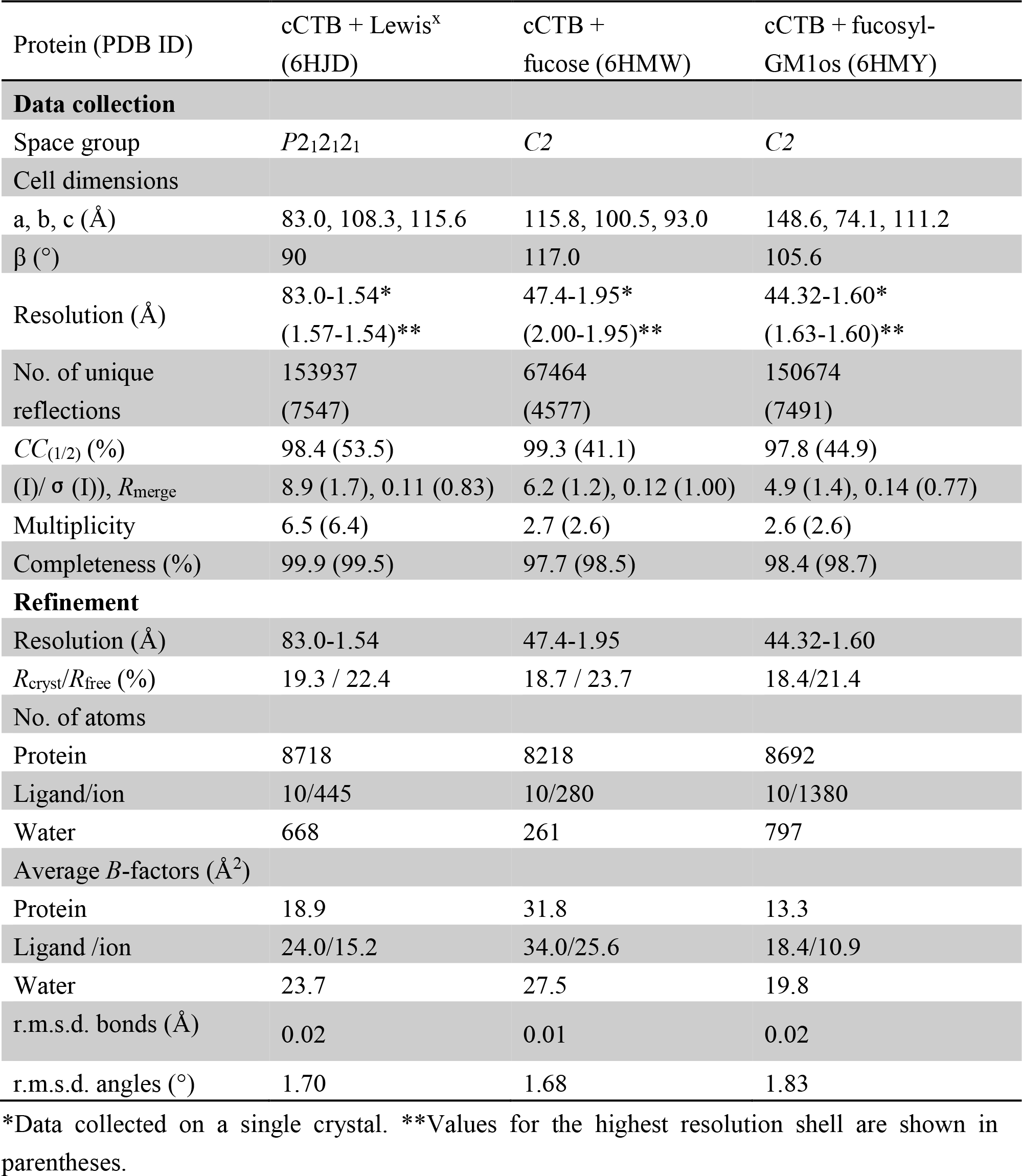
Data collection and refinement statistics.

| Protein (PDB ID) | cCTB + Lewis<sup>x</sup><br>(6HJD) | cCTB +<br>fucose (6HMW) | cCTB + fucosyl-GM1os (6HMY) |
| --- | --- | --- | --- |
| **Data collection** |  |  |  |
| Space group | *P*2<sub>1</sub>2<sub>1</sub>2<sub>1</sub> | *C*2 | *C*2 |
| Cell dimensions |  |  |  |
| a, b, c (Å) | 83.0, 108.3, 115.6 | 115.8, 100.5, 93.0 | 148.6, 74.1, 111.2 |
| β (°) | 90 | 117.0 | 105.6 |
| Resolution (Å) | 83.0-1.54*<br>(1.57-1.54)** | 47.4-1.95*<br>(2.00-1.95)** | 44.32-1.60*<br>(1.63-1.60)** |
| No. of unique<br>reflections | 153937<br>(7547) | 67464<br>(4577) | 150674<br>(7491) |
| *CC*<sub>(1/2)</sub> (%) | 98.4 (53.5) | 99.3 (41.1) | 97.8 (44.9) |
| (*I*)/ σ (*I*), *R*<sub>merge</sub> | 8.9 (1.7), 0.11 (0.83) | 6.2 (1.2), 0.12 (1.00) | 4.9 (1.4), 0.14 (0.77) |
| Multiplicity | 6.5 (6.4) | 2.7 (2.6) | 2.6 (2.6) |
| Completeness (%) | 99.9 (99.5) | 97.7 (98.5) | 98.4 (98.7) |
| **Refinement** |  |  |  |
| Resolution (Å) | 83.0-1.54 | 47.4-1.95 | 44.32-1.60 |
| *R*<sub>cryst</sub>/*R*<sub>free</sub> (%) | 19.3 / 22.4 | 18.7 / 23.7 | 18.4/21.4 |
| No. of atoms |  |  |  |
| Protein | 8718 | 8218 | 8692 |
| Ligand/ion | 10/445 | 10/280 | 10/1380 |
| Water | 668 | 261 | 797 |
| Average *B*-factors (Å<sup>2</sup>) |  |  |  |
| Protein | 18.9 | 31.8 | 13.3 |
| Ligand /ion | 24.0/15.2 | 34.0/25.6 | 18.4/10.9 |
| Water | 23.7 | 27.5 | 19.8 |
| r.m.s.d. bonds (Å) | 0.02 | 0.01 | 0.02 |
| r.m.s.d. angles (°) | 1.70 | 1.68 | 1.83 |

\*Data collected on a single crystal. \*\*Values for the highest resolution shell are shown in parentheses.

#### CTB in complex with L-fucose

Recent studies suggested that Le^x^ and similar structures might also bind to the GM1 binding site^32^. However, to date all relevant crystal structures of CTB contain fucosylated sugars in the secondary binding site^19, 20^. To alleviate concerns that protein purification by galactose affinity chromatography may hinder access to the GM1 binding site due to the presence of residual galactose (even though Gal is not observed in our crystal structures), we produced cCTB by TALON affinity chromatography exploiting its known affinity to Ni^2+^ or Co^2+^ ions^39^. To directly test the hypothesis by Cervin *et al.* that fucosylated sugars may bind to the GM1 binding site, TALON-purified cCTB was co-crystallized with L-fucose in a molar ratio of 1:500 (B-subunit to ligand). The crystals belong to space group *C2* and contained two CTB pentamers in the asymmetric unit. The structure was refined to 1.95 Å and an *R*_free_ value of 23.7% (Table 1). The electron density of the loop regions and the C-terminal Asn103 were more disordered compared to the cCTB-Le^x^ structure, however, overall the structure is well defined. Electron density for β-L-fucose is observed in nine of ten secondary binding sites (Figure 3 a,b, Supplementary Table S2). We did not observe electron density corresponding to L-fucose in the GM1 binding site, confirming that fucose only binds to the secondary binding site. Additionally, L-fucose was found sandwiched between two B-pentamers and covalently attached to some of the N-terminal residues. The latter is likely due to non-enzymatic glycosylation^40^. These interactions are unlikely to be biologically relevant and probably caused by the high molar excess of L-fucose added during crystallization.

**Figure 3.**
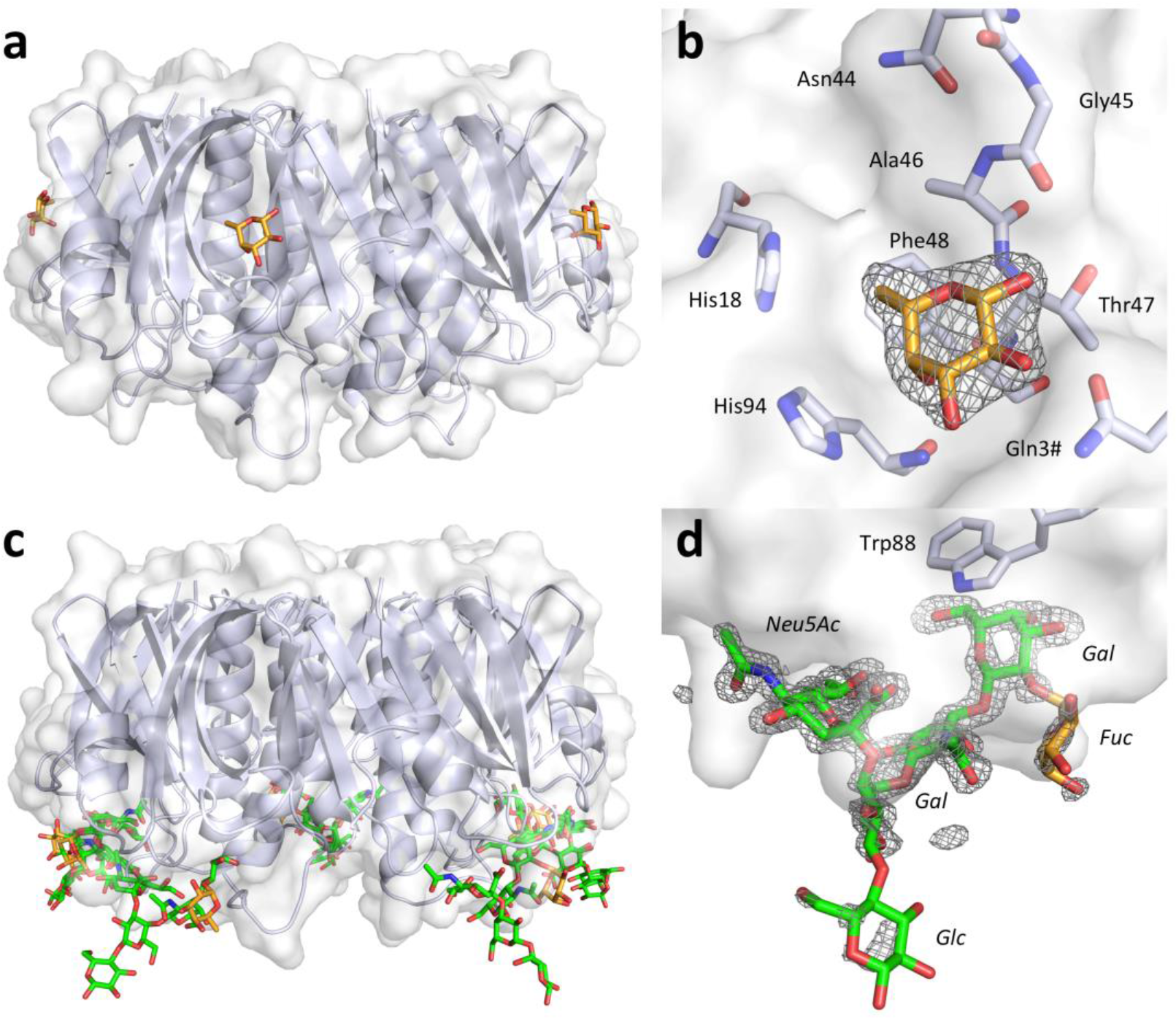
CTB complexes with L-fucose and fucosyl-GM1os. L-fucose binds to the secondary CT binding site, while fucosyl-GM1os binds to the primary binding site, facing the cell membrane. The cholera toxin B-pentamer is shown in cartoon (light blue) and transparent surface representations (white), and the ligands depicted in stick representation. (a) X-ray structure of cCTB in complex with L-fucose (orange sticks; PDB ID: 6HMW, this work); side view. (b) Close-up view of the secondary binding site, with σA-weighted F_o_-F_c_ electron density map for L-fucose (grey mesh, contoured at 3.0σ, generated before placing the ligand), and selected residues shown in stick representation, amino acid residues are labelled with 3-letter code. (c) X-ray structure of cCTB in complex with fucosyl-GM1os (green sticks, with the terminal fucose highlighted in orange; PDB ID: 6HMY, this work). (d) Close-up view of the primary binding site, with σA-weighted F_o_-F_c_ electron density map for fucosyl-GM1os (grey mesh, contoured at 3.0O, generated before placing the ligand) and Trp88 shown in stick representation. Carbohydrate residues are labelled in italics. The terminal fucose and glucose residues show weaker electron density compared to the four core residues of fucosyl-GM1os.

#### CTB in complex with fucosyl-GM1os

Our data strongly suggest that the toxin secondary binding site is the sole binding site for Le^x^ and similar sugars. There is, however, one fucosylated oligosaccharide that would be expected to bind to the primary binding site, namely fucosyl-GM1os. Fucosyl-GM1 binds CT almost as strongly as GM1^41^. We crystallized TALON-purified cCTB with fucosyl-GM1os in a 1:10 molar ratio (B-subunit to ligand), yielding crystals that diffracted to 1.6 Å resolution. The structure was refined to an .R_free_ value of 21.4%. Inspection of the electron density revealed that fucosyl-GM1os indeed binds to the primary binding site, similarly to GM1os, with the additional fucose residue facing outwards toward the solvent (Figure 3 c). The electron density for the fucose residue is less well defined compared to the other sugar residues (Figure 3 d), suggesting that it binds more weakly. It also has higher *B*-factors (average *B*-factors for sugar ring atoms in chain A: Fuc>GalNAc>Gal, 28.6>20.8>17.1 Å^2^). This explains the small difference in CT affinity between fucosyl-GM1 and GM1, as the fucose residue is not a major contributor to binding. In addition to the primary binding sites, we observe fucose binding in some of the secondary binding sites of cCTB. In these sites, fucosyl-GM1os is not well resolved, however, electron density extending from the fucose is compatible with a larger oligosaccharide like fucosyl-GM1os.

### Protein interaction by SPR

#### CTB and Le^x^ triaose

To determine the binding affinity of cCTB to Le^x^, we performed SPR experiments, in which the toxin B-pentamer was immobilized on the SPR chip and the sugar was injected as the analyte over a range of different concentrations. Le^x^ triaose was found to have a lower binding affinity to cCTB than Le^y^ tetraose or A-Le^y^ pentaose, which both feature a second fucose residue (*K*_d_=10 ± 3 mM (Le^x^ triaose), compared to 1.05 ± 0.04 mM (Le^y^) and 2.2 ± 0.1 mM (A-Le^y^)^19^; Supplementary Figure S1).

#### CT holotoxin and Le^x^ triaose and tetraose

Hitherto all crystal structures of CTB complexes with fucosylated receptors, including the structures reported here, suggest that Le^x^-related antigens bind exclusively to the secondary binding site of CT. To verify this binding mode in solution, we performed SPR experiments with holotoxin variants harbouring mutations in the primary binding site (W88K) or the BGA binding site (H18A, H18AH94A) (Figure 4, Table 2). Native folding of these variants was confirmed by circular dichroism (CD) spectroscopy (Supplementary Figure S2). Toxin variants or wild-type CT were immobilized on SPR chips, before injecting oligosaccharides at different concentrations. First, we showed that substitution of W88A in the primary binding site prevented GM1os binding (Figure 4). Since all other toxin variants bound GM1os, this confirmed an intact primary binding site. We then compared binding of Le^x^ oligosaccharides to wild-type CT and CT variants.

**Figure 4.**
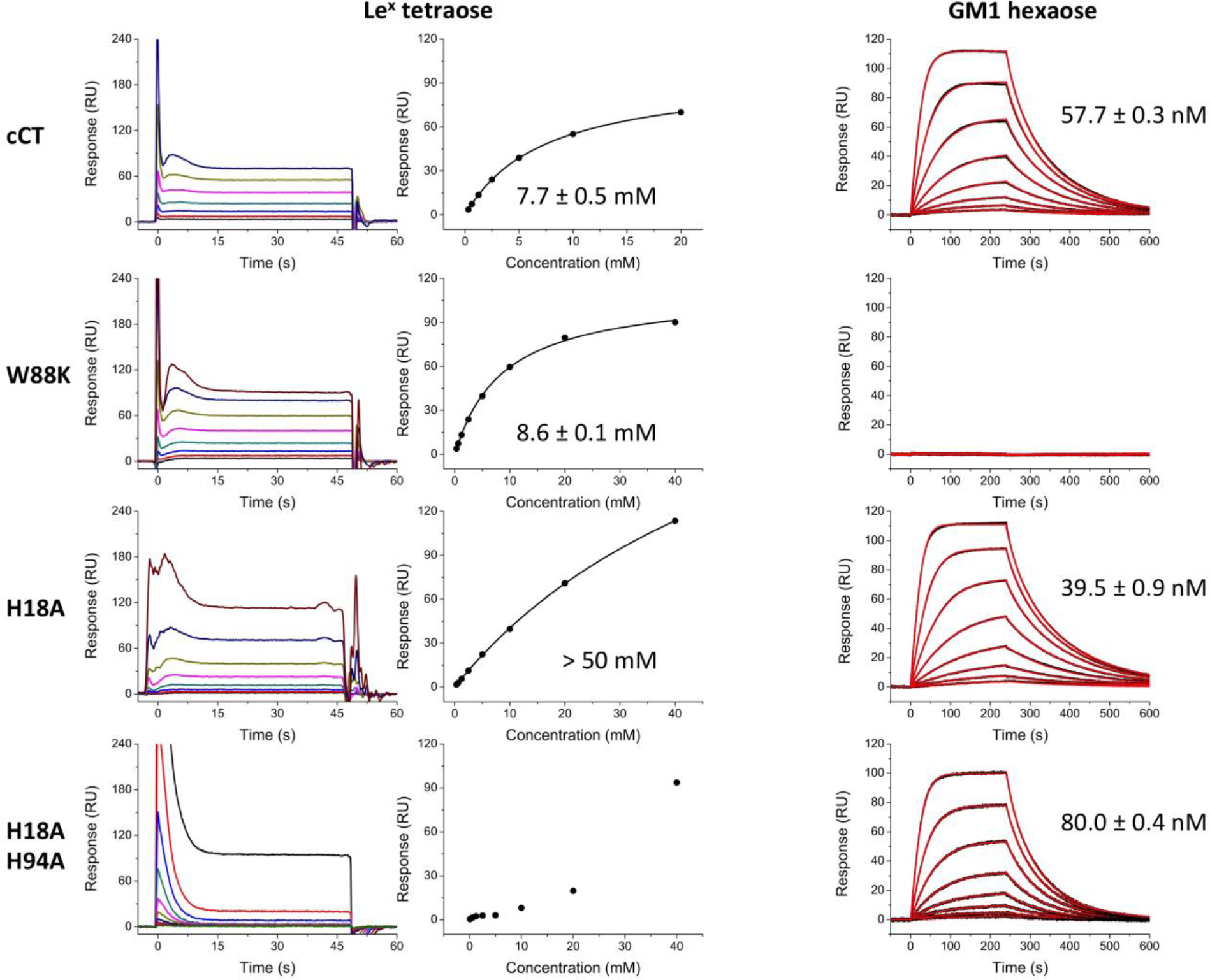
SPR sensorgrams and affinity plots for cholera toxin variants. SPR experiments were performed with cCT holotoxins coupled to the sensor chip and using Le^x^ tetraose (sensograms and corresponding plots of steady state response against concentration) or GM1os (fitted to 1:1 model) as analytes, as indicated in the panel legends. The *K*_d_ value for H18A + Le^x^ could not be calculated since saturation was not reached upon addition of 40 mM ligand. H18AH94A only showed weak interaction for the highest Le^x^ concentrations used; therefore, the affinity plot does not show a saturation-binding curve.

**Table 2.** *K*_d_ values for cCTB and CT variants and glycans GM1os, Le^x^ triaose and tetraose determined by SPR

|  | GM1os | Le<sup>x</sup> triaose | Le<sup>x</sup> tetraose |
| --- | --- | --- | --- |
| cCTB | not tested | $10 \pm 3$ mM | not tested |
| cCT | $57.7 \pm 0.3$ nM | $8 \pm 3$ mM | $7.7 \pm 0.5$ mM |
| W88K | no binding | not tested | $8.6 \pm 0.1$ mM |
| H18A | $39.5 \pm 0.9$ nM | not tested | > 50 mM |
| H18AH94A | $80.0 \pm 0.4$ nM | not tested | very weak binding |

Le^x^ triaose binds equally well to cCTB and cCT (*K*_d_=10 ± 3 mM for cCTB, 8 ± 3 mM for cCT) (Supplementary Figure S1). This result was expected, since there is no major structural difference between free CTB and the B-pentamer in the holotoxin^13, 42^. In our crystal structures, Le^x^ and related sugars pre-dominantly bind with the α-anomer of the reducing end GlcNAc, whereas relevant glycoconjugates on the cell surface are β-glycosidically linked to proteins or lipids. Le^x^ tetraose, with a fixed β-anomer, and Le^x^ triaose bound equally well to cCT (*K*_d_=8.1 ± 3.2 mM for triaose, 7.7 ± 0.5 mM for tetraose), suggesting that the triaose core is mainly responsible for binding and that the linkage does not significantly affect binding affinities.

Next, we determined binding affinities for all toxin variants using Le^x^ tetraose (Figure 4, Table 2). The primary binding site variant W88K was found to bind Le^x^ tetraose with an affinity comparable to wild-type CT (*K*_d_=8.6 ± 0.1 mM for W88K, 7.7 ± 0.5 mM for cCT), whereas H18A, which features a mutation in the secondary binding site, exhibited significantly reduced binding to Le^x^ tetraose (*K*_d_>50 mM for H18A, 7.7 ± 0.5 mM for cCT), in agreement with our structural data. This reduced binding affinity is not due to disruption of the primary binding site, since GM1os binding was found to be even stronger than for the wild-type protein (*K*_d_=39.5 ± 0.9 nM for H18A, 57.7 ± 0.3 nM for cCT; Figure 4). The double mutant H18AH94A, which was designed to disrupt the water network in the binding site and additionally precludes H-bond formation to the His94 side chain, only bound Le^x^ tetraose at the highest analyte concentration applied (40 mM), but bound to GM1os almost as strongly as wild-type CT (*K*_d_=80.0 ± 0.39 nM for H18AH94A, 57.7 ± 0.3 nM for cCT). Our results confirm Le^x^ binding to the secondary binding site of CT and suggest crosstalk between the primary and secondary binding sites, since mutations in the latter affect GM1 binding.

## Discussion

### Le^x^ and similar fucosylated sugars bind to the secondary binding site of CT

We set out to explore the molecular interaction of CT with fucosylated sugars, in particular Le^x^. Already twenty years ago, CTB was shown to bind fucose^43^. More recently, evidence was presented that BGAs may act as functional toxin receptors for the related LTB^44, 45^, and fucosylated glycan structures were shown to be functionally active and cause cellular uptake of CT^30,32,33^. For example, CTB binding to jejunal epithelial cells can be blocked by the Le^x^ trisaccharide or a monoclonal antibody against Le^x 32^ Crosslinking and immunoprecipitation of CTB-bound cellular proteins revealed that CTB binds to glycoproteins modified with Le^x 32^ It was, however, unclear how Le^x^ binds to CT. Also G33D, a CTB variant with greatly reduced affinity to GM1^15, 46^, showed weaker binding to Le^x 32^ Since Gly33 is located in the GM1 binding site, it was suggested that also Le^x^ might bind to the primary binding site^32^. To explore this possibility, Cervin *et al.* performed competition experiments with plate-bound GM1 or Le^x^ and CTB pre-incubated with the free sugars^32^. Competition experiments were also performed with human granulocytes, T84 and Colo205 cells. Despite the fact that these cell lines only express low levels of GM1, preincubation of CTB with GM1os had an inhibitory effect. Taken together, these results were interpreted to indicate that GM1 competitively blocks fucosylated sugars from binding to the CT primary binding site^32^. However, it should be noted that even at high concentrations, GM1os could not completely block CTB binding and Le^x^ could not block binding to plate-bound GM1.

Here we show that Le^x^ binds solely to the secondary binding site of cCTB, and even L-fucose alone (when co-crystallized in 500-fold molar excess) did not bind to the primary binding site. Moreover, SPR experiments with toxin variants confirmed that disruption of the secondary binding site, but not the primary binding site, lowered the affinity to Le^x^ (Figure 4, Table 2). We also found that Le^x^ bound more weakly to CTB than Le^y^ or A-Le^y^, which is in good agreement with inhibitor studies identifying Le^y^ as the most potent small-molecule inhibitor of CTB^33^. The binding mode of Le^x^ is highly similar to that of Le^y^ tetraose and A-Le^y^ pentaose^19^, which both feature an additional fucose residue (Figure 2 c). The partial competition of GM1 and Le^x^ observed by Cervin *et al?^1^* is most likely due to allosteric cross-talk, in line with results from a recent NMR study of the homologous LTB^47^. Similarly, we found that the CT secondary binding site variant H18A exhibits increased GM1os binding. Furthermore, even a small number of receptors in the cell-based assays could allow CT uptake, due to the high affinity of GM1 to CTB, which could explain the findings in the recent study^32^. However, even though Le^x^ and related fucosylated glycans and glycoconjugates do not bind to the primary CT receptor site, clearly additional fucose residues allow for additional attachment points that could interfere with toxin uptake, explaining the strong potency of a fucose-based polymer as CT inhibitor ^33^.

### Comparison of ligand structures to structure-activity relationship data

L-fucose has been shown to block CTB binding in cell culture, serving as a promising starting point for the design of novel cholera inhibitors^33^. Recently, Wands *et al.* described important molecular determinants of L-fucose by competition binding assays of CTB to various intestinal epithelial cells^33^. Their structure-activity relationship (SAR) studies showed that the stereochemical positioning of the terminal fucose is crucial for CTB binding. While the removal of hydroxyl groups on C3 or C4 (OH3, OH4) had a significant effect on CTB binding, substitution of the hydroxyl group at C2 (OH2) did not result in reduced binding (compare Figure 2 c). The addition of a hydroxyl group to C6 was tolerated, but the removal of C6 resulted in a complete loss of CTB binding suggesting the importance of a hydrophobic patch within the binding pocket. Addition of a methyl group to OH1, locking the anomeric hydroxyl group in the β-configuration, resulted in less efficient CTB binding compared to L-fucose or α-locked fucose. Finally, the SAR study showed increased binding of α1,2 linked fucose compared to α1,3 linked fucose^33^.

Our current ligand structures and previous structures of cCTB in complex with Le^y^ or A-Le^y^ (5ELB and 5ELD^19^) show that fucose OH3 can form one or two hydrogen bonds to His94 and to a conserved water molecule (not shown in Figure 2), explaining the observed reduced CTB binding upon its removal in the SAR study. OH4 was reported to contribute the strongest to CTB binding^33^, which is in good agreement with the fact that OH4 forms two hydrogen bonds to backbone atoms of residues 47 and 94 (Figure 2 d). The crystal structures also identify a match for the proposed key hydrophobic residues interacting with C6 in Phe48, possibly together with Ala46 (Figure 2 d).

cCTB binds L-fucose with its free anomeric hydroxyl group in the β-configuration (Figure 3 b), thus free fucose is not limited to bind in the α-configuration. However, locking the fucose in the β-configuration by the addition of a methyl group would cause steric clashes with residues Thr47 and Gly45, explaining why α-linked fucose bound stronger to CTB than β-linked fucose^33^.

According to Wands *et al.,* OH2 does not contribute significantly as a hydrogen bond acceptor^33^, whereas we observe a hydrogen bond to Gln3# from the adjacent B-subunit (Figure 2 d). However, their conclusion was based on a compound with a fluorine replacing the OH2 group (rather than with OH2 removed), and it is still under debate if fluorine can form hydrogen bonds^48, 49^.

Finally, Wands *et al.* found increased inhibition of CTB binding by fucosyllactose with α1,2-linked compared to α1,3-linked fucose residues^33^. The latter showed no difference compared to L-fucose alone. The crystal structure of CTB in complex with Le^y^ (5ELB^19^) clearly shows that both α1,2-linked and α1,3-linked fucose can bind to CT. However, the preferred orientation appears to be highly influenced by the precise details of ligands and toxins. For example, the structure of a CTB-LTB chimera in complex with A-HMO exhibited exclusive binding of Fucα2 in the fucose binding pocket of the toxin^20^.

In conclusion, the structural data presented here are a very good match to the reported structure-activity relationship data, pointing to the same binding site identified in both studies, *i.e.* the secondary binding site of the toxin.

### What are the roles of GM1 and fucosylated receptors? Are the binding sites connected by allosteric cross-talk?

A large body of research points to GM1 and fucosylated structures facilitating toxin binding and uptake^10,14–18,30,32,33^, however, little is known about the exact localization and expression of specific fucosylated structures on healthy intestinal cells and tissues^32,50–54^. It is not clear which fucosylated structures are the major cellular attachment sites besides GM1. However, all evidence points towards glycoproteins with type-2 core structures similar to Le^x^, Le^y^ and related BGAs^19,23–25,30,32,33^. In particular, Le^x^ has been shown to allow CTB binding to intestinal cells^32^, and Le^y^ is emerging as the most potent known small-molecule inhibitor of CTB, even superior to the GM1os in some cell lines^33^.

Whether the binding of GM1 and fucosylated structures happens simultaneously or sequentially is also unknown, nor what triggers or causes toxin uptake. Recent crystal structures from our lab suggest that CTB can bind sugars at both binding sites at the same time, specifically Le^y^ tetraose and galactose^19^. However, it has not yet been shown if this also holds true for GM1os. A recently published hetero-multivalent binding model suggests that CTB first binds to a high-affinity ligand such as GM1. Subsequent binding occurs much more readily (up to 10,000-fold faster) also for low-affinity ligands like GM2, since binding is then confined to the 2D membrane surface^55^. In fact, cooperativity was found to be enhanced for a heterogeneous mixture of ligands^55^. In a biological context, it seems more likely that the toxins first bind to the broadly available fucosylated structures on the cell surface or mucus layer until they can bind to the few available GM1 receptors, preventing the flow in the small intestine to remove unbound toxins from the intoxication site^19^. In addition, *V. cholerae* has other virulence factors that may influence receptor distribution and availability, *e.g.* neuraminidase VcN^56^. It has been shown very recently that VcN can remodel intestinal polysialylated gangliosides into GM1 to provide an increased numbers of cellular receptors for CT^56^, and it is well conceivable that fucosylated glycoconjugates serve as intermediate anchoring points until VcN unveils enough GM1 receptors.

Allosteric cross-talk between the two binding sites may facilitate toxin binding and uptake. GM1 binding to CTB is known to be positively cooperative^57–59^, and the small structural changes associated with cooperative binding to the primary site could easily be transmitted to the secondary binding site, for example *via* the helix connecting the two sites or by stabilization of the loop region comprising residues 55-60^60^. Recent NMR data for the homologous LTB from *E. coli* (ETEC) are consistent with such a cross-talk between binding sites^47^. Likewise, CT variant H18A showed increased GM1os affinity compared to wild-type CT, even though the mutation is in the secondary binding site. Cross-talk between the two sites could also explain the partial competition of GM1 and fucosylated receptors and the reduced binding of G33D to fucosylated structures observed by Cervin *et al.*^32^.

### Cholera blood group dependence: Why are secretors protected?

Many diseases show blood group association^61^, and cholera is the textbook example. Based on high-resolution structures of CTB in complex with BGAs, we recently elucidated the molecular basis of this phenomenon, *i.e.,* why individuals with blood group O experience the most severe symptoms^19^. Le^y^, characteristic for blood group O, exhibits a dual-binding mode and binds more strongly to CTB than the BGAs of blood groups A or B^19^, allowing for more efficient cellular uptake and more severe intoxication^31^.

Besides the commonly known ABO(H) blood group system, the secretor status of an individual can determine disease outcome. For example, ‘non-secretors’ are protected from Norwalk virus since suitable docking sites are not available at the infection site (reviewed by Heggelund *et al.*^61^). For cholera, however, it remains unclear why so-called ‘secretors’, who display BGAs in their small intestinal mucosa, are protected from severe cholera compared to ‘non-secretors’^62, 63^. Non-secretors have a non-functional *Se (FUT2)* gene, which codes for an α1,2-fucosyltransferase that is important for the generation of BGAs on epithelial and endothelial cells and most body fluids^64^. Non-secretors therefore display a distinct set of receptors on their intestinal cells that is enriched in the Lewis^a^ (Le^a^) and Le^x^ antigens, and characterized by limited fucosylation^52,64,65^. Interestingly, when probing jointly for secretor status and ABH(O) blood group, only secretors with blood group A or B showed less severe cholera symptoms, whereas the disease burden for blood group O individuals was equally high independent of their secretor status^63^, which could be explained if the H-antigen characteristic of blood group O itself is a risk factor for severe cholera.

The H-antigen exists in two types, type-1 and type-2, with different linkages, both of which contain an α1,2-linked fucose. Addition of a second fucose residue (Fucα4 or Fucα3) then gives rise to Le^b/y^ determinants, respectively. If the same fucose is added to H-antigen precursors instead, this yields Le^a/x^ epitopes. So far, all evidence points towards only type-2 antigens being able to bind to the CT^11,19,20^, focusing the attention on Le^y^ and Le^x^. The lower affinity of Le^x^ compared to the other BGAs makes it unlikely that Le^x^ binding is the cause for more efficient CT uptake in non-secretors. There are only limited data regarding the expression of fucosyltransferases and the distribution of glycoconjugates in human tissues^32,50–53,66^. Moreover, it is unknown which fucosylated CT receptors are relevant *in vivo.* The available data, however, suggest that compared to the villi, the deeper parts of the intestinal membranes are unaffected by the non-functional FUT2 of non-secretors, due to expression of an alternative 1,2-fucosyltransferase encoded by the *H* gene^65, 67^. Non-secretors cannot secrete soluble ABO(H) glycans, but might still present suitable docking sites *(i.e.,* Le^y^) on these deeper membrane regions, independent of their secretor status, and cooperative binding is expected to be enhanced for hetero-multivalent binding including GM1^55^. Indeed, Cervin *et al.* observed tighter binding of CTB to the crypts than to the villi^32^. Due to the lack of soluble BGAs, non-secretors would moreover lack competition by soluble and mucin-bound CT receptors, leading to more efficient CT uptake and more severe cholera symptoms.

### Conclusions and perspective

The importance of fucosylated sugars for cholera intoxication has recently been in the limelight^19,23–25,30–33^, but there has been limited structural information on their interaction with the CT. Here we show that fucosylated receptors, except fucosyl-GM1, bind solely to the secondary CT binding site, in contrast to recent suggestions^32^. We provide detailed information on how Le^x^ and L-fucose bind to CTB, and corroborate our findings with quantitative binding data of holotoxin variants. In addition, we present the first CT variant - H18A - with reduced affinity to fucosylated sugars, but increased affinity to GM1os. This not only lends further support to our hypothesis of a possible cross-talk between the primary and secondary CT binding sites^68^, but may also be of great practical value. Currently CT is widely used as a marker of GM1 and lipid rafts^69^. We suggest to replace wild-type CTB with variant H18A as more specific GM1 marker, to limit the number of false-positive results caused by the interaction with non-GM1 receptors. Furthermore, CT variants H18A and H18AH94A, being devoid of a functional secondary binding site, are predicted to facilitate the analysis of cellular uptake in cellular and organoid models^31, 70^ by allowing discrimination of the effects caused by interaction with primary and secondary sites. Finally, this study is expected to facilitate the design of more potent fucose-based CT inhibitors and CTB-containing vaccines, which may save countless human lives.

## Materials and Methods

### Mutagenesis

Vector pARCT5 contains an arabinose-inducible CT operon, with signal sequences derived from the LT-IIb B gene^71^. Site-directed mutagenesis was performed using the manufacturer’s protocol (Q5^®^ Site-Directed Mutagenesis Kit, NEB) and DNA oligos (Eurofins Genomics) shown in Table 3. Successful mutagenesis was verified by Sanger sequencing using primers Seq 1-3 (Eurofins Genomics, GATC).

**Table 3:**
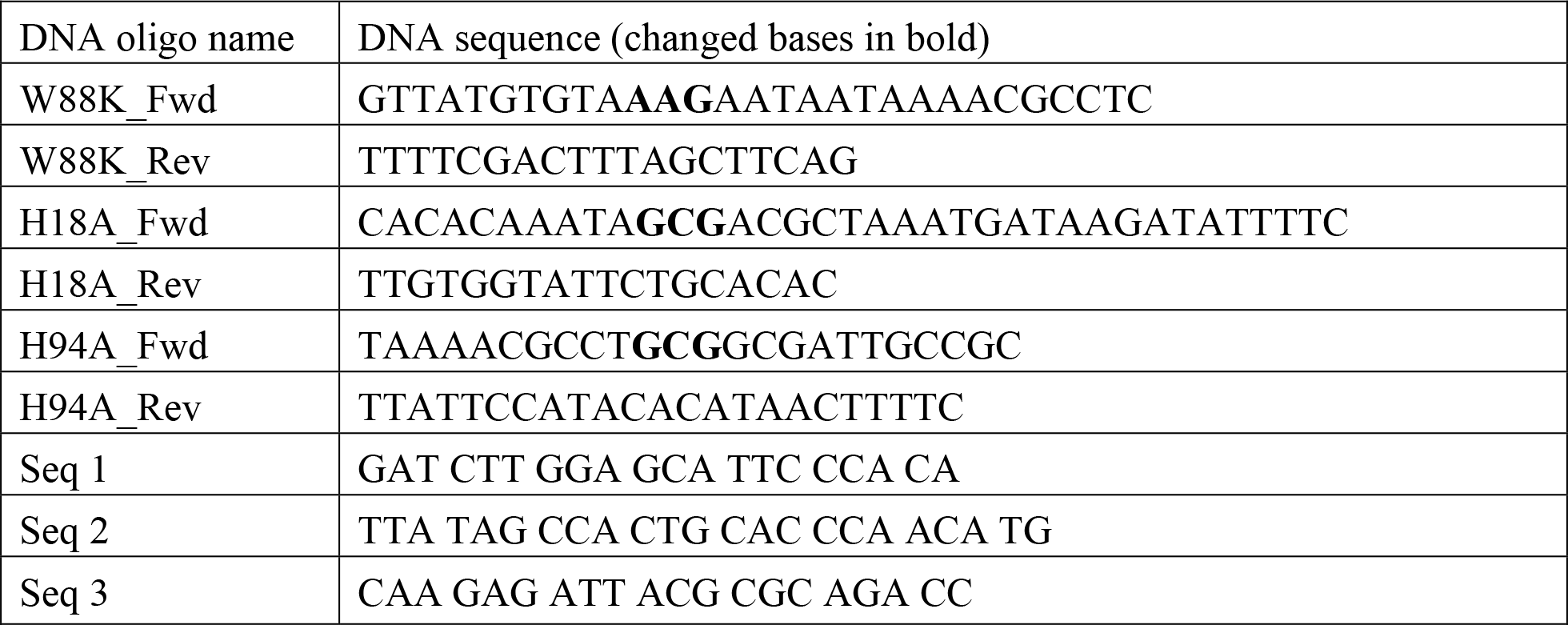
DNA oligonucleotides used in this study

### Expression of classical CTB

Protein expression was performed essentially as described previously^19^. Briefly, the gene for cCTB (Uniprot: Q57193) was heterologously expressed in *E. coli* BL21 (DE3) using a cCTB-pET21b+ construct. For protein production, cells were grown at 37°C in LB medium containing ampicillin until OD_600_ nm of 0.5 was reached. The temperature was reduced to 25°C and isopropyl-β-D-thiogalactopyranoside (IPTG) was added to a final concentration of 0.5 mM to start cCTB production. Cells were harvested after 14-18 h by centrifugation (6900 ⨯g, 20 min, 4°C) and the pellet was re-suspended in ice-cold sucrose buffer (20 mM Tris/HCl, 25% (w/v) sucrose, 5 mM EDTA at pH 8.0). After 15 min on ice, the solution was centrifuged (8000 ⨯g, 20 min, 4°C) and the pellet was re-suspended in periplasmic lysis buffer (5 mM MgCl_2_, 150 μg/mL lysozyme, DNase). The solution was kept cold for 30 min, centrifuged (8000 ⨯g, 20 min, 4°C) and dialyzed for several hours against PBS in a Snakeskin tube (Thermo Scientific, 3500 MWCO).

### Expression of CTB and CT holotoxins for TALON chromatography

The gene for cCTB (Uniprot: Q57193) was heterologously expressed in *E. coli* BL21 (DE3) using a cCTB-pET21b+ construct. For protein production, cells were grown at 37°C in LB medium containing ampicillin until OD_600_ nm of 0.5 was reached. The temperature was reduced to 25°C and IPTG was added to a final concentration of 0.5 mM to start cCTB production.

The genes for CT and CT variants (W88K, H18A, H18AH94A) were heterologously expressed in OverExpress^™^ C43 (DE3) cells (Sigma) using pARCT5 or pARCT5 derivatives. For protein production, cells were grown at 37°C in TB medium containing chloramphenicol until OD_600_ nm of 2.0 was reached. L-arabinose was added to a final concentration of 0.2% (w/v) to start holotoxin production.

Cells were harvested after 14-18 h (CTB) or 3 h (holotoxin) by centrifugation (6900 ⨯g, 20 min, 4°C) and the pellet was re-suspended in 1/60^th^ volume of TALON A buffer (50 mM sodium phosphate, 300 mM NaCl, pH 8) with 1 mg of polymyxin B (Sigma Aldrich) per mL, cOmplete^™^ Protease Inhibitor (Roche) and benzonase (EMD Millipore) and shaken at 37°C for 15 min. Insoluble debris and cells were removed from the periplasmic extracts by centrifugation (8000 xg, 20 min, 4°C). The filtered supernatant was directly applied to the TALON affinity column.

### Purification of cCTB using D-galactose affinity chromatography

The protein solution was loaded onto a D-galactose-sepharose affinity column (Thermo Scientific) and eluted using 300 mM galactose in PBS. Fractions containing pure cCTB were pooled and concentrated using Vivaspin 20 mL concentrator tubes (5000 MWCO, PES membrane, Sartorius). The protein was subjected to size-exclusion chromatography using PBS and a Superdex 75 column mounted on an ÄKTA purifier (GE Healthcare). Fractions with toxin were pooled, dialyzed o.n. against Tris buffer (20 mM Tris/HCl, 200 mM NaCl, pH 7.5), concentrated to 3-9 mg/mL, snap-frozen in liquid nitrogen and stored at −80°C.

### Purification of CTB and CT holotoxins using TALON affinity chromatography

As alternative to the D-galactose-sepharose affinity purification (to preclude contamination with galactose), the periplasmic extract of cCTB or CT (wild-type and variants) was loaded onto a preequilibrated HiTrap TALON crude (GE healthcare) and eluted with TALON B buffer (50 mM sodium phosphate, 300 mM NaCl, 50 mM imidazole at pH 8) in order to avoid residual D-galactose bound to the protein. The protein solution was concentrated using Vivaspin concentrator tubes (10000 MWCO, PES membrane, Sartorius) and loaded onto a Superdex 75 column mounted on an ÄKTA purifier (GE Healthcare) equilibrated with Tris buffer (20 mM Tris/HCl, 200 mM NaCl at pH 7.5) or PBS. Fractions with cCTB pentamer (Tris buffer) were pooled, concentrated to 3 mg/mL, snap-frozen in liquid nitrogen and stored at −80°C. Fractions with holotoxins (PBS) were pooled and stored at 4°C.

### CD spectroscopy and MS

Correct folding of all CT variants was verified by circular dichroism spectroscopy (J-810 CD spectrometer, JASCO). Prior to the measurement, buffer exchange to 10 mM sodium phosphate buffer pH 7.5 was performed using Vivaspin concentrator tubes. All CT variants were characterized by SDS-PAGE analysis, tryptic digestion and mass spectrometry (performed by the UiO core facility). The amino acid substitution H94A was only confirmed indirectly by DNA sequencing, since tryptic digestion and MS did not yield a suitable C-terminal peptide.

### Co-crystallization of CTB complexes

cCTB (Gal-affinity) and Le^x^ triaose (Gly049, Elicityl) or cCTB (TALON) and fucosyl-GM1os (GLY103, Elicityl) were mixed at a molar ratio of 1:10 (B-subunit to ligand). cCTB (TALON) and L-fucose (F2252, Sigma Aldrich) were mixed at a molar ratio of 1:500 (B-subunit to ligand) and all samples were incubated at RT for 2 h. Initially, crystals were obtained by sitting-drop vapour-diffusion using a crystallization robot (Oryx4, Douglas Instruments) at 20°C in the Morpheus screen condition A12. For CTB and Le^x^ triaose another crystal form was obtained in the Morpheus screen condition A4^72^. Crystals of condition A12 were optimized for all ligands by several rounds of hanging-drop vapour-diffusion experiments. Final crystals for the cCTB complex with Le^x^ were obtained by mixing protein-ligand solution (1 μL, 3.3 mg/mL) 1 : 1 with crystallization buffer (Tris (base)/BICINE pH 8.5, 8% PEG1000, 8% PEG3350, 8% MPD, 0.03 M MgCl_2_, 0.03 M CaCl_2_) and 0.2 μL seed solution (microseeding, Seed bead kit, Hampton research). cCTB-L-fucose co-crystals were obtained by mixing protein-ligand solution (1 μL, 3 mg/mL) 1:1 with crystallization buffer (Tris (base)/BICINE pH 8.7, 10% PEG1000, 10% PEG3350, 10% MPD, 0.03 M MgCl_2_, 0.03 M CaCl_2_). Final crystals for the cCTB complex with fucosyl-GM1os were obtained by mixing protein-ligand solution (1.5 μL, 1.5 mg/mL) 1:1 with crystallization buffer (Tris (base)/BICINE at pH 8.7, 6% PEG1000, 6% PEG3350, 6% MPD, 0.03 M MgCl_2_, 0.03 M CaCl_2_) and cryoprotection was achieved by transferring this crystal to crystallization buffer with higher PEG and MPD concentrations (Tris (base)/BICINE pH 8.7, 10% PEG1000, 10% PEG3350, 10% MPD, 0.03 M MgCl_2_, 0.03 M CaCl_2_. Crystals were harvested, mounted in loops, and flash-cooled in liquid nitrogen (cCTB-L-fucose, cCTB-fucosyl-GM1os) or flash-cooled in a nitrogen cryo-stream (cCTB-Le^x^).

### X-ray data collection and refinement

Synchrotron data collection for the cCTB complex with Le^x^ was performed at ID23-1, ESRF, Grenoble, France (100 K, 0.9770 Å). Data collection for cCTB complexes with L-fucose or fucosyl-GM1os was performed at BioMAX, Max IV, Lund, Sweden (100 K, 0.9795 Å and 0.980802 Å, respectively). Data were processed with XDS^73^ and NEGGIA/DECTRIS (cCTB-L-fucose, cCTB-fucosyl-GM1os) or *xia2* and *DIALS* (cCTB-Le^x^)^74, 75^ and *AIMLESS* from the *CCP4* software suite^76, 77^. Diffraction cut-offs were chosen based on the assessment of the anisotropic CC1/2 and the quality of the electron density (Table 1). Both crystal forms of cCTB-Le^x^ belong to space group *P*212121 and contain two B-pentamers in the asymmetric unit, but have different cell parameters. For the cCTB-Le^x^ model, we used the highest resolution data set (1.5 Å) from an optimized Morpheus A12 condition, which crystallized under similar conditions, and with the same space group and pentamer arrangement (“top-to-top”) as the cCTB complex with Le^y^ tetraose reported previously (5ELB^19^). cCTB-L-fucose and cCTB-fucosyl-GM1os crystals belong to space group *C2* and contain two B-pentamers in the asymmetric unit. The structures were solved by molecular replacement using *Phaser*^78^ from the CCP4 software suite^76, 77^ and search model 5ELB^19^, from which ligands and water molecules had been manually removed. To avoid potential model bias, five cycles of refinement including two cycles with simulated annealing (starting temperature of 5000 K) were carried out with the Phenix software suite^79^. The final model was obtained after several cycles of manual building with Coot^80^, followed by refinement with REFMAC5^81^. Initial refinement steps involved local NCS restraints, while final refinement steps involved TLS parameterization *(REFMAC5,* automatic, 5 cycles)^82^. Water molecules were placed using COOT:Find_waters and then manually inspected for several criteria, including distances from hydrogen-bond donors / acceptors and quality of the electron-density. Most of the disulfide bridges are partly reduced, due to minor radiation damage. Le^x^ triaose, GM1os and fucosyl-GM1os were built using MAKE LIGAND (AceDRG)^83^ from the CCP4 software suite^76, 77^ and isomeric SMILES strings. The restraints for a Thr-Fuc bond were generated using JLigand^84^. Le^x^ triaose, fucosyl-GM1os and α-L-fucose or β-L-fucose were included last to avoid model bias. To improve the density for the terminal fucose residue, GM1os was included prior to fucosyl-GM1os. For the cCTB complex with fucose, additional elongated electron density was found in two of the primary binding sites, however, the origin of the density could not be identified, even with Polder^85^ maps calculated with the Phenix software suite^79^ (the density was clearly not compatible with a sugar ring). PDB_REDO^86^ was used to evaluate the models before final refinement steps. Occupancies were refined by evaluating the difference Fourier maps and by comparing the *B*-factors of the ligands with interacting protein atoms. The final models were analysed using the Analyse geometry task of the *CCP*4 software suite^76, 77^. The percentages of amino acid residues occupying the favoured, allowed and outlier regions in the Ramachandran plot are 97.5/2.5/0.0% for cCTB-Le^x^, 97.4/2.4/0.2% for cCTB-L-fucose, and 97.7/2.3/0.0% for cCTB-fucosyl-GM1os, respectively. Figures were generated using PyMol (Schrödinger LLC), α-helices and β-strands were assigned using STRIDE^87^.

### Surface plasmon resonance spectroscopy

SPR analyses were performed using Series S CM5 sensor chips and a Biacore T100 biosensor system (Biacore Life Sciences, GE Healthcare, Uppsala, Sweden). Due to the high cost of the oligosaccharides combined with the relatively low affinity typical for carbohydrate-protein interactions, experiments were performed in duplicates or triplicates. Proteins were immobilized on the chip as described previously^19^. Le^x^ tetraose (GLY050, Elicityl) and triaose were dissolved in running buffer (10 mM HEPES/NaOH, 140 mM NaCl, 3 mM EDTA, 0.005 % P20, pH 7.4) and injected in concentrations from 78.125 μM up to 40 mM for 45 s. Dissociation was monitored for additional 15 s. The regeneration was not needed for Le^x^ tetraose, whereas for Le^x^ triaose a short pulse (6 s) of 3 mM NaOH was injected to completely remove it from the surface. Binding of 3.125-200 nM GM1os (GM1a, GLY096, Elicityl) was monitored for 240 s, with additional 420 s for dissociation. The regeneration of the chip was the same as for Le^x^ triaose. All binding steps were performed at 30 μL/min and 25°C. The data were evaluated using Biacore T100 Evaluation software. For GM1os, data were fitted to the Langmuir 1:1 interaction model. Other binding affinities were determined from plots of steady-state analyte binding levels against the concentration.

## Data availability

The coordinates and structure factors have been deposited with the Protein Data Bank with accession codes: 6HJD, 6HMW and 6HMY.

## Acknowledgements

We are grateful to our colleagues Per Eugen Kristiansen for help with CD spectroscopy, Bernd Thiede for MS measurements and Kaare Bjerregaard-Andersen for synchrotron data collection for cCTB-fucosyl-GM1 crystals. We would further like to thank Professor Randall K. Holmes for their generous gift of plasmid pARCT5, and the staff at the ESRF and Max IV for assistance and support using beamlines ID23-1 and BioMAX, respectively. This work was funded by the University of Oslo (position of J.B.H.) and the Norwegian Research Council (grant 247730). Further support was by iNEXT and the Norwegian Graduate School in Biocatalysis (BioCat).

## Author Contributions

U.K. conceived the research; J.B.H. designed the experiments; J.B.H. and V.H. performed the experiments; J.B.H., J.E.H., V.H, G.A. and U.K. analysed the data; U.K. validated the crystal structures, J.B.H. and U.K. wrote the manuscript, with input from all authors.

## Competing Financial Interests

The author(s) declare no competing financial interests.

## References

1 Sack, D. A., Sack, R. B., Nair, G. B. & Siddique, A. K. Cholera. Lancet 363, 223–233 (2004).

2 WHO. Yemen, Annual report 2017, https://reliefweb.int/report/yemen/world-health-organization-yemen-annual-report-2017 (Date of access:29/09/2018) (2018).

3 The Lancet Gastroenterology & Hepatology. Health catastrophe: the toll of cholera in Yemen. Lancet Gastroenterol Hepatol 2, 619, doi:10.1016/S2468-1253(17)30224-8 (2017).

4 Heggelund, J. E., Bjørnestad, V. A. & Krengel, U. Vibrio cholerae and Escherichia coli heat-labile enterotoxins and beyond. In The Comprehensive Sourcebook of Bacterial Protein Toxins (eds J. Alouf, D. Landant & M. R. Popoff) 195–229 (Elsevier, 2015).

5 Merritt, E. A. & Hol, W. G. J. AB_5_ toxins. Curr Opin Struct Biol 5, 165–171, doi:10.1016/0959-440X(95)80071-9 (1995).

6 Chinnapen, D. J., Chinnapen, H., Saslowsky, D. & Lencer, W. I. Rafting with cholera toxin: endocytosis and trafficking from plasma membrane to ER. FEMS Microbiol Lett 266, 129–137, doi:10.1111/j.1574-6968.2006.00545.x (2007).

7 Gill, D. M. & Meren, R. ADP-ribosylation of membrane proteins catalyzed by cholera toxin: basis of the activation of adenylate cyclase. Proc Natl Acad Sci U S A 75, 3050–3054 (1978).

8 Field, M., Rao, M. C. & Chang, E. B. Intestinal electrolyte transport and diarrheal disease (1). N Engl J Med 321, 800–806, doi:10.1056/NEJM198909213211206 (1989).

9 Holmgren, J. Comparison of the tissue receptors for Vibrio cholerae and Escherichia coli enterotoxins by means of gangliosides and natural cholera toxoid. Infect Immun 8, 851–859 (1973).

10 Holmgren, J., Lönnroth, I., Månsson, J. & Svennerholm, L. Interaction of cholera toxin and membrane GM1 ganglioside of small intestine. Proc Natl Acad Sci U S A 72, 2520–2524 (1975).

11 Ångström, J. et al. Novel carbohydrate binding site recognizing blood group A and B determinants in a hybrid of cholera toxin and Escherichia coli heat-labile enterotoxin B-subunits. J Biol Chem 275, 3231–3238 (2000).

12 Turnbull, W. B., Precious, B. L. & Homans, S. W. Dissecting the cholera toxin-ganglioside GM1 interaction by isothermal titration calorimetry. J Am Chem Soc 126, 1047–1054, doi:10.1021/ja0378207 (2004).

13 Merritt, E. A. et al. Crystal structure of cholera toxin B-pentamer bound to receptor GM1 pentasaccharide. Protein Sci 3, 166–175, doi:10.1002/pro.5560030202 (1994).

14 Fishman, P. H., Moss, J. & Vaughan, M. Uptake and metabolism of gangliosides in transformed mouse fibroblasts. Relationship of ganglioside structure to choleragen response. J Biol Chem 251, 4490–4494 (1976).

15 Jobling, M. G. & Holmes, R. K. Analysis of structure and function of the B subunit of cholera toxin by the use of site-directed mutagenesis. Mol Microbiol 5, 1755–1767 (1991).

16 Wolf, A. A. et al. Ganglioside structure dictates signal transduction by cholera toxin and association with caveolae-like membrane domains in polarized epithelia. J Cell Biol 141, 917–927 (1998)

17 Badizadegan, K. et al. Floating cholera toxin into epithelial cells: functional association with caveolae-like detergent-insoluble membrane microdomains. Int J Med Microbiol 290, 403–408, doi:10.1016/S1438-4221(00)80052-1 (2000).

18 Fujinaga, Y. et al. Gangliosides that associate with lipid rafts mediate transport of cholera and related toxins from the plasma membrane to endoplasmic reticulm. Mol Biol Cell 14, 4783–4793, doi:10.1091/mbc.e03-06-0354 (2003).

19 Heggelund, J. E. et al. High-resolution crystal structures elucidate the molecular basis of cholera blood group dependence. PLoS Pathog 12, e1005567, doi:10.1371/journal.ppat.1005567 (2016).

20 Holmner, Å. et al. Novel binding site identified in a hybrid between cholera toxin and heat-labile enterotoxin: 1.9 Å crystal structure reveals the details. Structure 12, 1655–1667, doi:10.1016/j.str.2004.06.022 (2004). Erratum in: Structure 15, 253 (2007).

21 Nair, G. B. et al. Cholera due to altered El Tor strains of Vibrio cholerae O1 in Bangladesh. J Clin Microbiol 44, 4211–4213, doi:10.1128/JCM.01304-06 (2006).

22 Mekalanos, J. J. et al. Cholera toxin genes: nucleotide sequence, deletion analysis and vaccine development. Nature 306, 551–557 (1983).

23 Heggelund, J. E. et al. Both El Tor and classical cholera toxin bind blood group determinants. Biochem Biophys Res Commun 418, 731–735, doi:10.1016/j.bbrc.2012.01.089 (2012).

24 Mandal, P. K. et al. Towards a structural basis for the relationship between blood group and the severity of El Tor cholera. Angew Chem Int Ed Engl 51, 5143–5146, doi:10.1002/anie.201109068 (2012).

25 Vasile, F. et al. Comprehensive analysis of blood group antigen binding to classical and El Tor cholera toxin B-pentamers by NMR. Glycobiology 24, 766–778, doi:10.1093/glycob/cwu040 (2014).

26 Morita, A., Tsao, D. & Kim, Y. S. Identification of cholera toxin binding glycoproteins in rat intestinal microvillus membranes. J Biol Chem 255, 2549–2553 (1980).

27 Monferran, C. G., Roth, G. A. & Cumar, F. A. Inhibition of cholera toxin binding to membrane receptors by pig gastric mucin-derived glycopeptides: differential effect depending on the ABO blood group antigenic determinants. Infect Immun 58, 3966–3972 (1990).

28 Balanzino, L. E., Barra, J. L., Galvan, E. M., Roth, G. A. & Monferran, C. G. Interaction of cholera toxin and Escherichia coli heat-labile enterotoxin with glycoconjugates from rabbit intestinal brush border membranes: relationship with ABH blood group determinants. Mol Cell Biochem 194, 53–62, doi:10.1023/A:1006971913175 (1999).

29 Holmner, Å., Askarieh, G., Ökvist, M. & Krengel, U. Blood group antigen recognition by Escherichia coli heat-labile enterotoxin. J Mol Biol 371, 754–764, doi:10.1016/j.jmb.2007.05.064 (2007).

30 Wands, A. M. et al. Fucosylation and protein glycosylation create functional receptors for cholera toxin. Elife 4, e09545, doi:10.7554/eLife.09545 (2015).

31 Kuhlmann, F. M. et al. Blood group O-dependent cellular responses to cholera toxin: parallel clinical and epidemiological links to severe cholera. Am J Trop Med Hyg 95, 440–443, doi:10.4269/ajtmh.16-0161 (2016).

32 Cervin, J. et al. GM1 ganglioside-independent intoxication by cholera toxin. PLoS Pathog 14, e1006862, doi:10.1371/journal.ppat.1006862 (2018).

33 Wands, A. M. et al. Fucosylated molecules competitively interfere with cholera toxin binding to host cells. ACS Infect Dis, doi:10.1021/acsinfecdis.7b00085 (2018).

34 Glass, R. I. et al. Predisposition for cholera of individuals with O blood group. Possible evolutionary significance. Am J Epidemiol 121, 791–796 (1985).

35 Swerdlow, D. L. et al. Severe life-threatening cholera associated with blood group O in Peru: implications for the Latin American epidemic. J Infect Dis 170, 468–472 (1994).

36 Harris, J. B. et al. Blood group, immunity, and risk of infection with Vibrio cholerae in an area of endemicity. Infect Immun 73, 7422–7427, doi:10.1128/IAI.73.11.7422-7427.2005 (2005).

37 Harris, J. B. et al. Susceptibility to Vibrio cholerae infection in a cohort of household contacts of patients with cholera in Bangladesh. PLoS Negl Trop Dis 2, e221, doi:10.1371/journal.pntd.0000221 (2008).

38 Breimer, M. E., Hansson, G. C., Karlsson, K. A., Larson, G. & Leffler, H. Glycosphingolipid composition of epithelial cells isolated along the villus axis of small intestine of a single human individual. Glycobiology 22, 1721–1730, doi:10.1093/glycob/cws115 (2012).

39 Dertzbaugh, M. T. & Cox, L. M. The affinity of cholera toxin for Ni^2+^ ion. Protein Eng 11, 577–581 (1998).

40 Furth, A. J. Methods for assaying nonenzymatic glycosylation. Anal Biochem 175, 347–360, doi:10.1016/0003-2697(88)90558-1 (1988).

41 Masserini, M., Freire, E., Palestini, P., Calappi, E. & Tettamanti, G. Fuc-GM1 ganglioside mimics the receptor function of GM1 for cholera toxin. Biochemistry 31, 2422–2426, doi:10.1021/bi00123a030 (1992).

42 O’Neal, C. J., Amaya, E. I., Jobling, M. G., Holmes, R. K. & Hol, W. G. Crystal structures of an intrinsically active cholera toxin mutant yield insight into the toxin activation mechanism. Biochemistry 43, 3772–3782, doi:10.1021/bi0360152 (2004).

43 Mertz, J. A., McCann, J. A. & Picking, W. D. Fluorescence analysis of galactose, lactose, and fucose interaction with the cholera toxin B subunit. Biochem Biophys Res Commun 226, 140–144, doi:10.1006/bbrc.1996.1323 (1996).

44 Galvan, E. M., Diema, C. D., Roth, G. A. & Monferran, C. G. Ability of blood group A-active glycosphingolipids to act as Escherichia coli heat-labile enterotoxin receptors in HT-29 cells. J Infect Dis 189, 1556–1564, doi:10.1086/383349 (2004).

45 Galvan, E. M., Roth, G. A. & Monferran, C. G. Functional interaction of Escherichia coli heat-labile enterotoxin with blood group A-active glycoconjugates from differentiated HT29 cells. FEBS J 273, 3444–3453, doi:10.1111/j.1742-4658.2006.05368.x (2006).

46 Merritt, E. A. et al. Structural studies of receptor binding by cholera toxin mutants. Protein Sci 6, 1516–1528, doi:10.1002/pro.5560060716 (1997).

47 Hatlem, D., Heggelund, J. E., Burschowsky, D., Krengel, U. & Kristiansen, P. E. ^1^H, ^13^C, ^15^N backbone assignment of the human heat-labile enterotoxin B-pentamer and chemical shift mapping of neolactotetraose binding. Biomol NMR Assign 11, 99–104, doi:10.1007/s12104-017-9728-9 (2017).

48 Howard, J. A. K., Hoy, V. J., O’Hagan, D. & Smith, G. T. How good is fluorine as a hydrogen bond acceptor? Tetrahedron 52, 12613–12622, doi:10.1016/0040-4020(96)00749-1 (1996).

49 Dalvit, C., Invernizzi, C. & Vulpetti, A. Fluorine as a hydrogen-bond acceptor: experimental evidence and computational calculations. Chem. Eur. J. 20, 11058–11068, doi:10.1002/chem.201402858 (2014).

50 Björk, S., Breimer, M. E., Hansson, G. C., Karlsson, K. A. & Leffler, H. Structures of blood group glycosphingolipids of human small intestine. A relation between the expression of fucolipids of epithelial cells and the ABO, Le and Se phenotype of the donor. J Biol Chem 262, 6758–6765 (1987).

51 Finne, J. et al. Novel polyfucosylated N-linked glycopeptides with blood group A, H, X, and Y determinants from human small intestinal epithelial cells. J Biol Chem 264, 5720–5735 (1989).

52 Ravn, V. & Dabelsteen, E. Tissue distribution of histo-blood group antigens. APMIS 108, 1–28, doi:10.1034/j.1600-0463.2000.d01-1.x (2000).

53 Henry, S. M. Molecular diversity in the biosynthesis of GI tract glycoconjugates. A blood-group-related chart of microorganism receptors. Transfus Clin Biol 8, 226–230, doi:10.1016/S1246-7820(01)00112-4 (2001).

54 Robbe, C. et al. Evidence of regio-specific glycosylation in human intestinal mucins: presence of an acidic gradient along the intestinal tract. J Biol Chem 278, 46337–46348, doi:10.1074/jbc.M302529200 (2003).

55 Krishnan, P. et al. Hetero-multivalent binding of cholera toxin subunit B with glycolipid mixtures. Colloids Surf B Biointerfaces 160, 281–288, doi:10.1016/j.colsurfb.2017.09.035 (2017).

56 Alisson-Silva, F. et al. Human evolutionary loss of epithelial Neu5Gc expression and species-specific susceptibility to cholera. PLoS Pathog 14, e1007133, doi:10.1371/journal.ppat.1007133 (2018).

57 Schafer, D. E. & Thakur, A. K. Quantitative description of the binding of GM1 oligosaccharide by cholera enterotoxin. Cell Biophys 4, 25–40, doi:10.1007/BF02788553 (1982).

58 Lin, H., Kitova, E. N. & Klassen, J. S. Measuring positive cooperativity using the direct ESI-MS assay. Cholera toxin B subunit homopentamer binding to GM1 pentasaccharide. J Am Soc Mass Spectr 25, 104–110, doi:10.1007/s13361-013-0751-5 (2014).

59 Worstell, N. C., Krishnan, P., Weatherston, J. D. & Wu, H. J. Binding cooperativity matters: A GM1-like ganglioside-cholera toxin B subunit binding study using a nanocube-based lipid bilayer array. PLoS One 11, e0153265, doi:10.1371/journal.pone.0153265 (2016).

60 Holmner, Å. et al. Crystal structures exploring the origins of the broader specificity of Escherichia coli heat-labile enterotoxin compared to cholera toxin. J Mol Biol 406, 387–402, doi:10.1016/j.jmb.2010.11.060 (2011).

61 Heggelund, J. E., Varrot, A., Imberty, A. & Krengel, U. Histo-blood group antigens as mediators of infections. Curr Opin Struct Biol 44, 190–200, doi:10.1016/j.sbi.2017.04.001 (2017).

62 Chaudhuri, A. & DasAdhikary, C. R. Possible role of blood-group secretory substances in the aetiology of cholera. Trans R Soc Trop Med Hyg 72, 664–665 (1978).

63 Arifuzzaman, M. et al. Individuals with Le(a+b-) blood group have increased susceptibility to symptomatic Vibrio cholerae O1 infection. PLoS Negl Trop Dis 5, e1413, doi:10.1371/journal.pntd.0001413 (2011).

64 Henry, S., Oriol, R. & Samuelsson, B. Lewis histo-blood group system and associated secretory phenotypes. Vox Sang 69, 166–182, doi:10.1111/j.1423-0410.1995.tb02591.x (1995).

65 Glynn, L. E., Holborow, E. J. & Johnson, G. D. The distribution of blood-group substances in human gastric and duodenal mucosa. Lancet 273, 1083–1088 (1957).

66 Torrado, J., Blasco, E., Cosme, A., Gutierrez-Hoyos, A. & Arenas, J. I. Expression of type 1 and type 2 blood group-related antigens in normal and neoplastic gastric mucosa. Am J Clin Pathol 91, 249–254 (1989).

67 Mollicone, R., Bara, J., Le Pendu, J. & Oriol, R. Immunohistologic pattern of type 1 (Lea, Leb) and type 2 (X, Y, H) blood group-related antigens in the human pyloric and duodenal mucosae. Lab Invest 53, 219–227 (1985).

68 Holmner-Rocklöv, Å. Molecular recognition of carbohydrates-structural and functional characterisation of bacterial toxins and fungal lectins, PhD thesis, Chalmers University of Technology, Gothenburg, Sweden, (2005).

69 Chiricozzi, E., Mauri, L., Ciampa, M. G., Prinetti, A. & Sonnino, S. On the use of cholera toxin. Glycoconj J, doi:10.1007/s10719-018-9818-7 (2018).

70 Zomer-van Ommen, D. D. et al. Functional characterization of cholera toxin inhibitors using human intestinal organoids. J Med Chem 59, 6968–6972, doi:10.1021/acs.jmedchem.6b00770 (2016).

71 Jobling, M. G., Palmer, L. M., Erbe, J. L. & Holmes, R. K. Construction and characterization of versatile cloning vectors for efficient delivery of native foreign proteins to the periplasm of Escherichia coli. Plasmid 38, 158–173, doi:10.1006/plas.1997.1309 (1997).

72 Gorrec, F. The MORPHEUS protein crystallization screen. J Appl Crystallogr 42, 1035–1042, doi:10.1107/S0021889809042022 (2009).

73 Kabsch, W. Xds. Acta Crystallogr D Biol Crystallogr 66, 125–132, doi:10.1107/S0907444909047337 (2010).

74 Parkhurst, J. M. et al. Robust background modelling in DIALS. J Appl Crystallogr 49, 1912–1921, doi:10.1107/S1600576716013595 (2016).

75 Winter, G. et al. DIALS: implementation and evaluation of a new integration package. Acta Crystallogr D Struct Biol 74, 85–97, doi:10.1107/S2059798317017235 (2018).

76 Collaborative Computational Project, N.The CCP 4 suite: programs for protein crystallography. Acta Crystallogr D Biol Crystallogr 50, 760–763,doi:10.1107/S0907444994003112 (1994).

77 Potterton, L. et al. CCP4i2: the new graphical user interface to the CCP4 program suite. Acta Crystallogr D Struct Biol 74, 68–84, doi:10.1107/S2059798317016035 (2018).

78 McCoy, A. J. et al. Phaser crystallographic software. J Appl Crystallogr 40, 658–674, doi:10.1107/S0021889807021206 (2007).

79 Adams, P. D. et al. The Phenix software for automated determination of macromolecular structures. Methods 55, 94–106, doi:10.1016/j.ymeth.2011.07.005 (2011).

80 Emsley, P., Lohkamp, B., Scott, W. G. & Cowtan, K. Features and development of Coot. Acta Crystallogr D Biol Crystallogr 66, 486–501, doi:10.1107/S0907444910007493 (2010).

81 Murshudov, G. N. et al. REFMAC5 for the refinement of macromolecular crystal structures. Acta Crystallogr D Biol Crystallogr 67, 355–367, doi:10.1107/S0907444911001314 (2011).

82 Winn, M. D., Isupov, M. N. & Murshudov, G. N. Use of TLS parameters to model anisotropic displacements in macromolecular refinement. Acta Crystallogr D Biol Crystallogr 57, 122–133 (2001).

83 Long, F. et al. AceDRG: a stereochemical description generator for ligands. Acta Crystallogr D Struct Biol 73, 112–122, doi:10.1107/S2059798317000067 (2017).

84 Lebedev, A. A. et al. JLigand: a graphical tool for the CCP4 template-restraint library. Acta Crystallogr D Biol Crystallogr 68, 431–440, doi:10.1107/S090744491200251X (2012).

85 Liebschner, D. et al. Polder maps: improving OMIT maps by excluding bulk solvent. Acta Crystallogr D Struct Biol 73, 148–157, doi:10.1107/S2059798316018210 (2017).

86 Joosten, R. P., Long, F., Murshudov, G. N. & Perrakis, A. The PDB_REDO server for macromolecular structure model optimization. IUCrJ 1, 213–220, doi:10.1107/S2052252514009324 (2014).

87 Heinig, M. & Frishman, D. STRIDE: a web server for secondary structure assignment from known atomic coordinates of proteins. Nucleic Acids Res 32, W500–502, doi:10.1093/nar/gkh429 (2004).

